# VineColD: an integrative database for global historical tracing and real-time monitoring of grapevine cold hardiness

**DOI:** 10.1101/2025.01.07.631810

**Authors:** Hongrui Wang, Jason P. Londo

## Abstract

Cold hardiness is a crucial physiological parameter that determines the survival of grapevines during the dormant season. Accurate modeling and large-scale prediction of grapevine cold hardiness are essential for assessing the potential geographic distribution of grapevine cultivation, quantifying the impact of climate change on grapevine habitats, and ensuring the sustainability of the grape and wine industries in cool climate regions worldwide. However, until now, no comprehensive database has been available. In this research, we combined advanced automated machine learning techniques with extensive historical and current weather data to create an integrative database for grapevine cold hardiness: VineColD (https://cornell-tree-fruit-physiology.shinyapps.io/VineColD/). We developed the NYUS.2.1 model, an automated machine learning-based system for predicting grapevine cold hardiness and in this study, applied it to global historical weather data from 17,985 curated weather stations spanning 30° to 55° in both hemispheres from 1960 to 2024, resulting in the development of an integrative grapevine cold hardiness database and monitoring system. VineColD integrates both a global historical dataset and a daily updated regional cold hardiness system, offering a comprehensive resource to study grape cold hardiness for 54 grapevine cultivars. The platform provides multiple download options, from single-station data to complete datasets, and the interactive multi-functional R Shiny application facilitates data analysis and visualization. VineColD delivers critical insights into the impact of climate change on grapevine cultivation and supports a range of analytical functions, making it a valuable tool for grape growers and researchers.

## Introduction

The distribution of woody perennials across the world is predominantly influenced by climatic factors, with temperature extremes in both summer and winter playing a crucial role in determining their survival [1,2]. Climate change, characterized by increased frequency of unusual temperature extremes during the growing season or the dormant season, poses a threat to these species, including economically important perennial crops such as grapevine [3–5]. In cool or cold climate viticulture regions established closed to the temperature boundaries, extreme low temperatures in winter that exceed grapevine cold hardiness and late spring frosts that damage fragile fruitful tissues after budbreak are the leading causes of crop loss, affecting the ongoing viability and potential expansion of the grape and wine industries in these regions [6–8]. The accurate prediction of dormant season physiology, like bud cold hardiness and budbreak timing, can be used as guidance for the cultivation of grapevines in changing winters under climate change.

Our current understanding of grapevine dormant season physiology is based on the U-shape cold hardiness dynamic backboned by the theorical changing cold acclimation-deacclimation equilibrium in response to chilling accumulation and the response to daily temperature variation [9–17]. While the concurrent bud cold hardiness determines if grapevines could survive the extreme cold events in winter, bud cold hardiness is also a crucial physiological character effecting early spring phenology, including timing of budbreak [18]. The modelling of grapevine dormant physiology is therefore centered on the prediction of bud cold hardiness. Currently, the benchmark models for grapevine cold hardiness prediction includes WAUS.1, WAUS.2, WIUS.2, NYUS.1, NYUS.2 and an unnamed recurrent neural network (RNN)-based model. The WAUS.1 and its successor WAUS.2 were developed in Washington, US. These models are mechanistic models where continuous changes in bud cold hardiness, estimated as the lethal temperature that kills 50% of bud population (LT50), are arbitrarily phased with incremental time steps based on daily maximum and minimum temperature and cultivar-specific parameters [9,19]. WIUS.1 is derived from the modelling method of WAUS.2 but is trained using more the LT50 data of the interspecific cold hardy cultivars grown in Wisconsin, US [20]. The NYUS.1 model, a recent mechanistic model with empirically derived biological parameters, has been developed to incorporate a phased integration of cold acclimation and deacclimation responses based on the latest insights into the dynamics of dormancy-related cold hardiness in grapevines [21]. The RNN-based model, developed in Washington, US, is a deep learning model that uses unstructured time series weather data to predict the whole season bud LT50. The model was trained through multi-task learning, which leverages data across multiple grape cultivars to improve prediction accuracy, especially for the cultivars with limited training data [22]. Beyond the modeling technique and the restricted range of cultivars available for cold hardiness prediction, a notable limitation of all these models is that they were developed using data from a single location, potentially causing them to become excessively tailored to specific local conditions. For example, the underperformance of WAUS.1 and WAUS.2 has be reported when applying these models in other locations with different climate conditions [21,23,24]. The final model, NYUS.2, was developed using automated machine-learning (Auto-ML) and trained with 1,0157 cold hardiness data from nine locations and 72 features extracted from air temperature (engineered temperature parameters such as chilling and heat accumulation). This model has demonstrated exceptional robustness and outperformed the previous mechanistic models in the prediction of the cold hardiness of 45 cultivars in various viticultural regions in North America [24]. In addition to leveraging available hardware for enhanced computational speed through the Auto-ML engine, the high site-transferability of this model enables large-scale and multi-location tracing and forecasting of grapevine bud cold hardiness/cold damage, which is essential for the global assessment of the impact of climate change on grapevine dormant season physiology.

In this project, we coupled NYUS.2.1, an updated NYUS.2 model, with GHCNd (Global Historical Climatology Network daily) to calculate grapevine cold hardiness and potential cold damage of 54 grapevine cultivars at all accessible weather stations within the 30° – 55° viticultural zone of both hemispheres over the 64 dormant seasons from 1960 to 2024. Furthermore, we have created a real-time monitoring system that offers daily updates on grapevine cold hardiness for the major cool climate viticultural regions in North America. By combining various sources of grapevine cold hardiness prediction data, actual measurements with previous and current weather records, we developed an integrative database named VineColD (grape**Vine Col**d hardiness **D**atabase, https://cornell-tree-fruit-physiology.shinyapps.io/VineColD/). VineColD is now accessible through a web-based interactive platform that supports diverse functionalities like data analysis, visualization, as well as uploading and downloading of data, thereby providing a comprehensive tool for grape growers and researchers to navigate and adapt to the challenges posed by climate change.

## Methods and data collection

To develop an integrated database for the global historical tracing and regional real-time monitoring of grapevine cold hardiness/cold damage, we dynamically applied NYUS.2.1 on historical and current temperature records from the weather stations included in GHCNd. The entire process of model generation, data curation, model prediction and data analysis are briefed in Figure 1.

**Figure 1.**
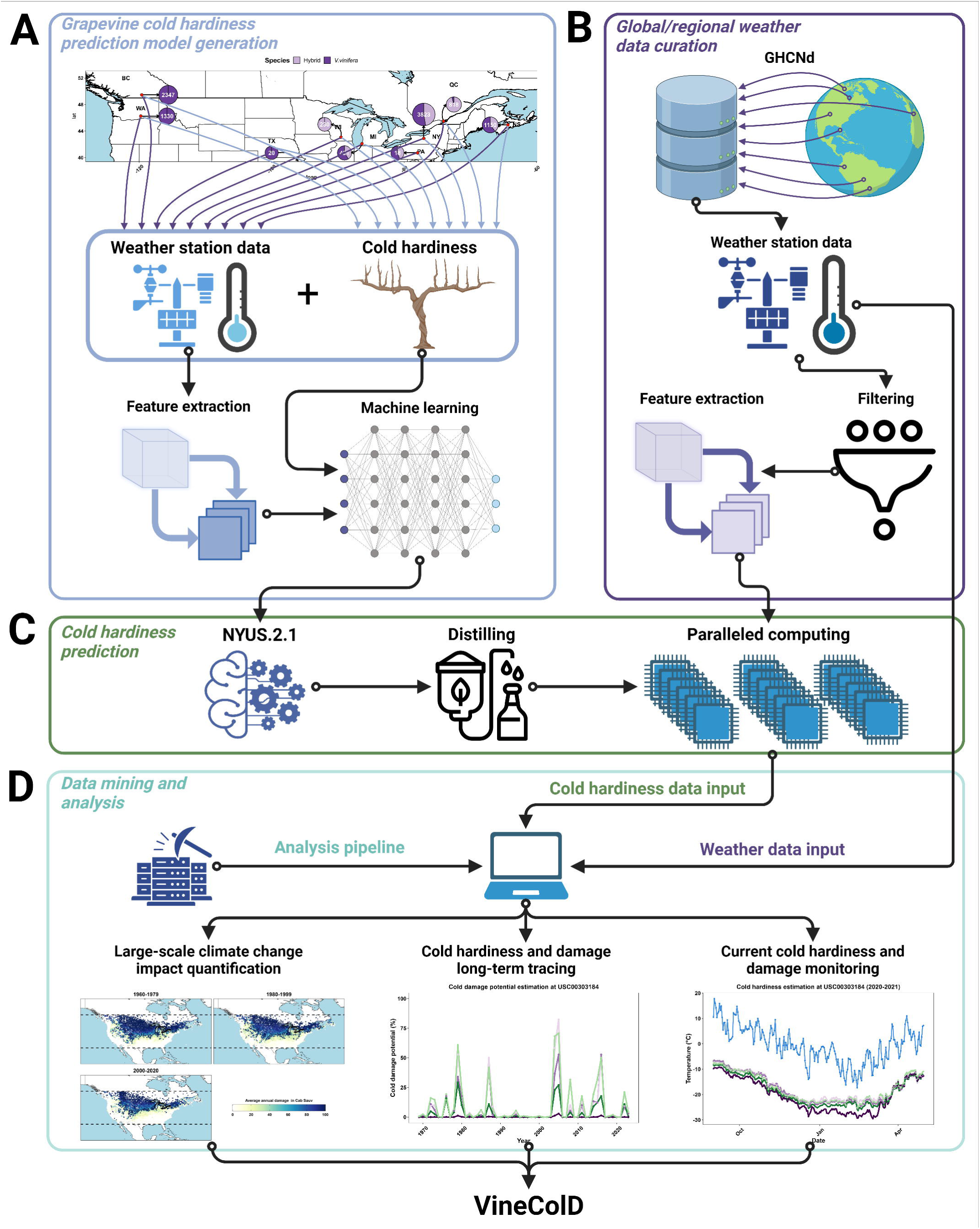
Schematic representation of the development of VineColD. A) Generation of the new grapevine cold hardiness prediction model, NYUS.2.1; B) Curation of global and regional weather data; C) Computation of grapevine cold hardiness prediction; D) Data processing and analysis of predicted grapevine cold hardiness for the construction of VineColD.

### Grapevine cold hardiness prediction model NYUS.2.1

The model used in this study for predicting grapevine cold hardiness, NYUS.2.1, is an updated iteration of the original NYUS.2, an automated machine learning-based model [24]. NYUS.2 was developed using on-site grapevine cold hardiness measurements collected from nine regions across North America between 2003 and 2022 (n == 10157), coupled with features derived from daily temperature data from local weather stations (Figure 1A). The model was trained using the AutoGluon automated machine learning engine (version 0.7.0) [25]. NYUS.2.1 builds on this foundation by incorporating the original training data along with additional cold hardiness measurements from three locations in New York State during the 2022-2023 and 2023-2024 dormant seasons (n == 11277). The training, testing and feature importance analysis were performed through the same pipeline of NYUS.2 by an upgraded version of AutoGluon (version 1.1.0) [24]. Model performance was evaluated using root mean square error (RMSE) by comparing predicted cold hardiness with observed cold hardiness. The NYUS.2.1 model offers improved predictive capacity for an expanded range of cultivars, increasing from 45 to 54. The prediction relies on the one-hot encoded cultivar information and features extracted from daily maximum and minimum temperatures only.

### Curation of global and regional temperature data

Two types of data curation were conducted to support grapevine cold hardiness predictions, each serving a different purpose: global data curation for historical tracing and regional data curation for real-time monitoring. The data curation process is summarized in Figure 1B.

To trace historical grapevine cold hardiness, temperature data from weather stations were curated using the Global Historical Climatology Network-Daily (GHCNd) (https://www.ncei.noaa.gov/products/land-based-station/global-historical-climatology-network-daily). To ensure accurate and efficient predictions while optimizing computational resources, a four-step filtering process was applied to the GHCNd dataset: record type filtering, geographic filtering, temporal filtering, and data quality filtering.

1. Record Type Filtering: Weather stations lacking continuous records of both daily maximum and minimum temperatures were excluded, as the features for NYUS.2.1 predictions are derived from daily temperature data [24]. Additionally, stations with less than two years of data were filtered out to ensure data stability.
2. Geographic Filtering: Since grapevines are predominantly cultivated between latitudes 30° and 50° [8,26,27], we filtered out weather stations outside the 30° to 55° latitude band. This range covers both current and potential future viticultural areas where cold stress may be a concern, considering the expected migration of grape and wine industries to cooler regions due to climate change [5,28–30].
3. Temporal Filtering: Stations that ceased recording before 1960 were excluded, along with any data before 1960-01-01, due to issues with accuracy and missing data.
4. Data Quality Filtering: Temperature records from each remaining weather station were segmented by dormant season (September 01 to May 01 for the northern hemisphere and March 01 to November 01 for the southern hemisphere). The dormant seasons with fewer than 220 days of temperature record were filtered out to ensure data quality for feature extraction.

The monitoring of current grapevine cold hardiness focused on the viticultural regionals in North America where cold damage poses a significant concern [6]. Temperature data for the current season was collected daily from onsite measurements at weather stations curated within the GHCNd dataset with a focus on the cool or cold climate viticultural regions in North America. Specifically, data was gathered from all GHCNd weather stations located within the geographic boundaries of North Dakota (NW), Maine (NE), Virginia (SE), Colorado (SW), and the states of Washington, Oregon, New Mexico and Texas in the U.S., as well as British Columbia, Québec, Ontario and Nova Scotia in Canada. No filtering was applied to these weather stations or their data. This daily data collection supports the near real-time updating of grapevine cold hardiness prediction. Additionally, temperature data forecast for the current season was collected from open-meteo (https://github.com/open-meteo/open-meteo) for the forecast of grapevine cold hardiness for the upcoming seven days.

### Feature extraction and grapevine cold hardiness prediction

The feature extraction for temperature data in NYUS.2.1 followed the same methodology as in NYUS.2 [24]. Hourly temperature was estimated based on daily maximum and minimum temperature using ‘stack_hourly_temps’ of the R package ‘chillR’ [31]. Features extracted from hourly temperature data were categorized into four groups: daily temperature descriptors (four features), cumulative temperature descriptors (seven features), exponential weighted moving average temperatures (30 features), and reverse exponential weighted moving average temperatures (30 features). Combined with 54 Boolean-type cultivar features and days in season (number of days after September 1), a total of 126 features were used for NYUS.2.1 predictions. The computation process for grapevine cold hardiness is outlined in Figure 1C. For historical grapevine cold hardiness tracing, NYUS.2.1 was distilled for model prediction using ‘distill = True’ option to manage computational resource demands [25]. Predictions were conducted on all the cultivars included in NYUS.2.1: 21 *Vitis. vinifera* cultivars and 33 *Vitis. hybrid* cultivars. We plan to establish an annual update for historical grapevine cold hardiness predictions to ensure the database remains current and accurate. For current grapevine cold hardiness monitoring, the full model was deployed, with daily predictions based on continuously updated features. These predictions also covered all cultivars in NYUS.2.1. In addition to NYUS.2.1, two previous benchmark models, NYUS.1 and WAUS.2 were also used for the predictions for ‘Cabernet Sauvignon’, ‘Concord’ and ‘Riesling’ [19,21].

For both purposes, feature extraction and model prediction for NYUS.2.1 was conducted with paralleled computation by distributing the main task to the available threads through ‘parallel’ and AutoGluon built-in parallelization, respectively [25]. The predicted grapevine cold hardiness of a cultivar is expressed as LT50, representing the temperature that kills 50% of the bud population [32].

### Cold damage estimation

Potential grapevine cold damage was estimated by comparing the predicted cold hardiness with the minimum temperature of the day. The potential (0-100) that a cold damage has occurred is estimated using a symmetric sigmoid function assuming that 10% and 90% of the potential that a damage has occurred when minimum temperatures is 2 °C above or below the predicted cold hardiness [24].

### Data processing and visualization

The processing, analysis and visualization of predicted grapevine cold hardiness/cold damage data are briefed in Figure 1D. For global historical grapevine cold hardiness tracing, predictions from NYUS.2.1 and the estimated cold damage data were organized by the weather station. At each curated station, the maximum cold hardiness and maximum cold damage per dormant season were calculated to visualize the overall impact of climate change over the 64 years between 1960 to 2024. Additionally, the data was segmented by individual seasons to visualize daily cultivar-specific cold hardiness dynamics in response to minimum temperature. For regional real-time grapevine cold hardiness monitoring, daily updated predictions were visualized across all curated weather stations. Geospatial analysis and visualization were conducted using R package ‘sp’ [33].

### VineColD

All the data generated from global historical cold hardiness tracing and regional real-time cold hardiness monitoring are combined as an integrative database, named VineColD (grape**Vine Col**d hardiness **D**atabase). VineColD is currently hosted in an R shiny-based application at https://cornell-tree-fruit-physiology.shinyapps.io/VineColD/ [34]. The global historical cold hardiness dataset is divided by weather station and is stored as .RDS in Box Cloud. Data analysis, visualization, and downloading are facilitated through API pull requests using the built-in function in the application based on ‘boxr’ R package [35]. Real-time cold hardiness monitoring is hosted in a separate R shiny-based application, which is mirrored in the VineColD application. All applications built for VineColD feature interactive figures created using the ‘plotly’ R package, interactive weather station maps generated with the ‘leaflet’ R package, and interactive tables made using the ‘DT’ R package [36–38].

## Results and discussion

### Performance of NYUS.2.1 in predicting grapevine cold hardiness

During model training, a total of 67 models were generated using various algorithms across different stacking levels (Figure 2A). The final model for NYUS.2.1, was selected as a “WeightedEnsembleL6,” a computationally optimized ensemble model developed after five levels of stacking. This model is composed of eight base or stacker models from stacking levels one and two. This model was chosen for its superior accuracy combined with higher computational efficiency, demonstrated by its relatively low inference latency (Figure 2A). Feature importance analysis using SHAP values [39] revealed that the top 15 most influential features included seven accumulative temperature descriptors: three chilling models (Utah, NC, and CU models) and four growing degree hours (GDH) with base temperatures of 0 °C, 4 °C, 7 °C, and 10 °C (Figure 2B). In model testing with unseen data (10% of the entire dataset, n == 1127), NYUS.2.1 demonstrated performance ranging from RMSE == 0.82 °C in the ‘BC’ sub-dataset to RMSE == 1.58 °C in the ‘Geneva, NY’ sub-dataset (Figure 2C). These prediction errors are minimum as compared to the prediction errors in the utilization/testing of the benchmark models [19,21,23,40]. The grapevine cold hardiness predictions by NYUS.2.1 exhibited a balanced distribution around the zero-error slope, which indicates minimal bias towards overestimation or underestimation (Figure 2C). This balanced error profile suggests an absence of systemic error, which is crucial for the large-scale monitoring of grapevine cold hardiness both locally and globally.

**Figure 2.**
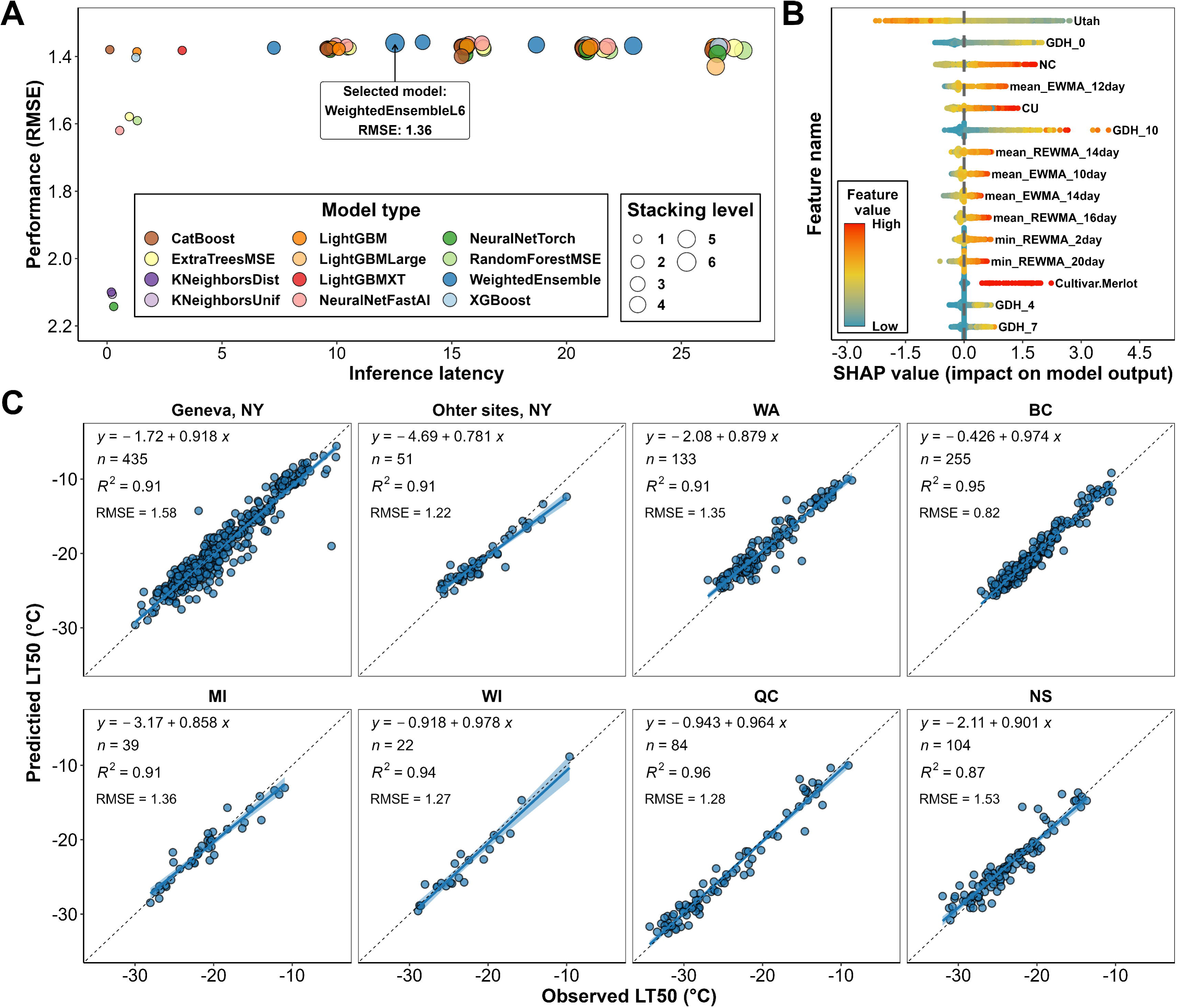
Model selection, feature importance analysis and model testing for the NYUS.2.1 model. A) The performance (RMSE) of all models generated during the NYUS.2.1 training process; B) Distribution of SHAP value for the top 15 features in the final model, with feature importance ranked by the mean absolute SHAP values across all testing samples; C) Performance of NYUS.2.1 in predicting cold hardiness across various sub-datasets within the testing data.

### Curation of temperature data

The geographical distribution and completeness of the historical records for the curated weather stations in the VineColD database are detailed in Figure 3. For global historical grapevine cold hardiness tracing, a total of 17,985 weather stations were curated after the filtering process. The geographical distribution of these stations is shown in Figure 3A, where each point represents a weather station. These stations are in 63 countries across six continents (Africa, Asia, Europe, North America, South America and Oceania). Over the 64 years from 1960 to 2024, the number of stations has steadily increased, reaching a peak in the early 2010s before experiencing a decline (Figure 3B). The detailed breakdown of weather station numbers by country reveals a significant contribution from the U.S. and Canada (Figure 3B). The average total record length is 31.8 years, with a notable subset of stations providing data for over six decades, which adds a significant temporal depth to the database (Figure 3C). The average number of missing years in the records is 4.8 years, and the histogram of missing years shows a scale-free distribution, indicating that most curated stations have maintained comprehensive records with minimal annual gaps (Figure 3D).

**Figure 3.**
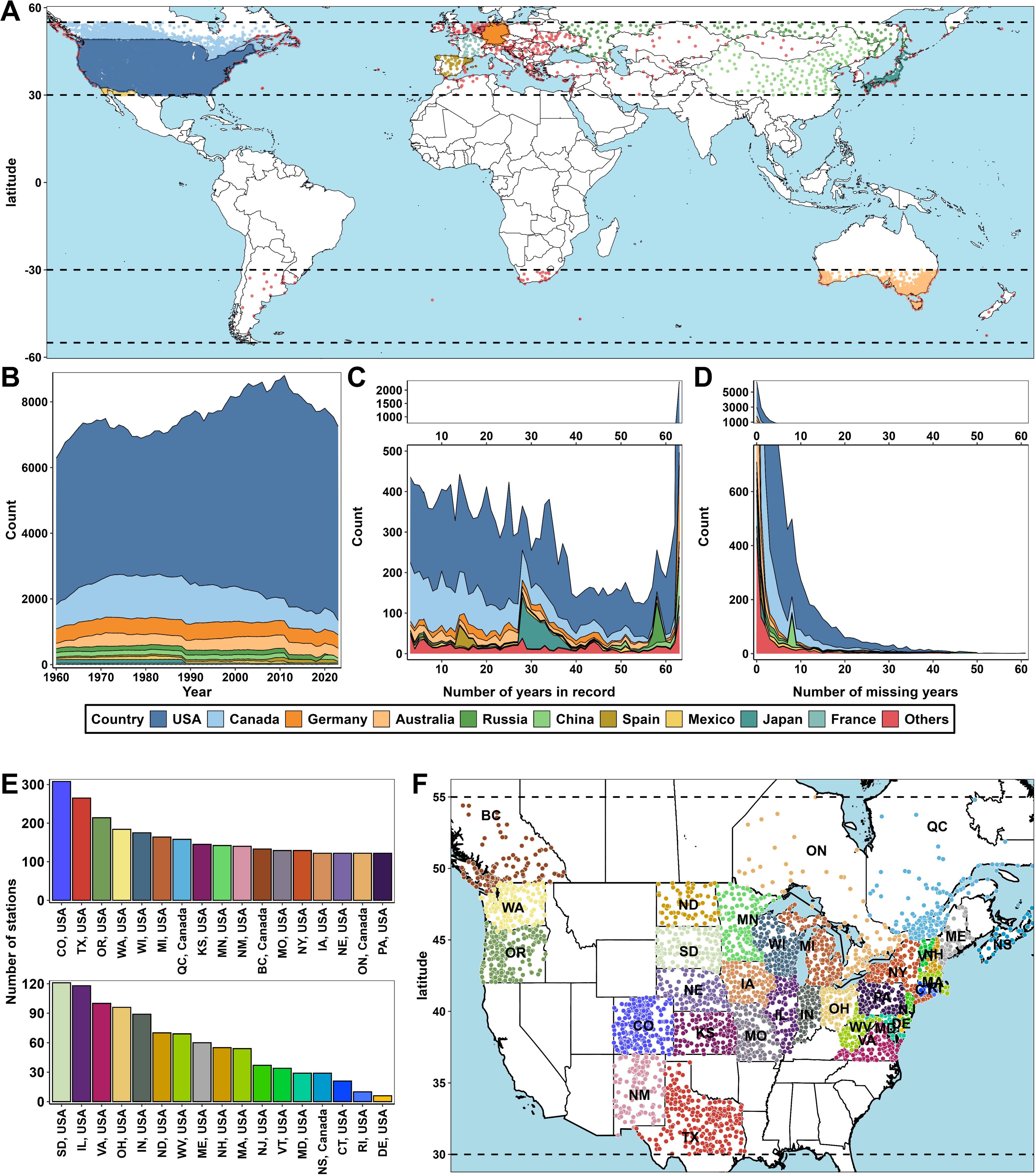
Curation of temperature data for VineColD. A) Distribution of weather stations in the global data curation process for historical grapevine cold hardiness tracing; B) Number of stations per year; C) Number of years recorded per station; D) Number of missing years per station; E) Number of stations per state/province in the regional data curation for real-time monitoring of grapevine cold hardiness in North America; F) Distribution of weather stations of the regional curation in North America.

The regional real-time grapevine cold hardiness monitoring system was launched in the early 2022-2023 dormant season and has since provided cold hardiness predictions across three dormant seasons. In the current system, a total of 3,772 weather stations have been curated, evenly distributed across the monitored regions (Figure 3E and F). Colorado (CO), Texas (TX), Oregon (OR), Washington (WA) and Wisconsin (WI) are the top five regions contributing the most weather stations (Figure 3E).

Overall, the extensive curation of weather stations in VineColD ensures a historically rich and geographically comprehensive dataset, providing a robust foundation for accurate and reliable grapevine cold hardiness modeling and climate impact studies.

### Functions of different components in VineColD

The predicted cold hardiness, estimated cold damage, historical temperature data and current temperature data were combined and used as the data input for VineColD. VineColD is deployed in a user interface that provides various data acquisition options and data visualizations methods towards the curation and interactive analysis of global historical and regional real-time grapevine cold hardiness (Figure 4A). The major components of VineColD application are ‘Historical data’, ‘Current data’ and ‘Archived data’. ‘Historical data’ supports three functions: ‘Station data’ allows users to select individual weather stations on a station map, download selected cold hardiness/damage data, and visualize seasonal maximum cold hardiness, maximum cold damage summaries, and detailed cold hardiness/cold damage dynamics for each season (Figure 4B); ‘Bulk download’ enables users to download cold hardiness data from grouped weather stations based on various filtering criteria such as start year, end year, missing year, continent, and country (Figure 4C); ‘North America long-term data viewer’ provides visualizations of average minimum temperatures, average maximum predicted cold hardiness, and average maximum estimated cold damage for all curated weather stations in North America between 1960 and 2024, along with corresponding station maps showing their geographic distribution (Figure 4D). ‘Current data’ hosts a daily-updated application for the current grapevine cold hardiness monitoring with predicted cold hardiness from three models, NYUS.2.1, NYUS.1 and WAUS.2, along with uploaded on-site measurements in the cool climate viticultural regions in the U.S (Figure 4E). The system automatically begins updating each year on September 15. The application enables the near real-time monitoring of grapevine cold hardiness and includes a ‘Data upload’ portal for users to upload onsite cold hardiness measurements, which are displayed and available for download the day after uploading. ‘Archived data’ hosts old versions of the ‘Current data’ application from the 2022-2023 and the 2023-2024 dormant seasons, including all the predicted and uploaded measurement data (Figure 4F and G).

**Figure 4.**
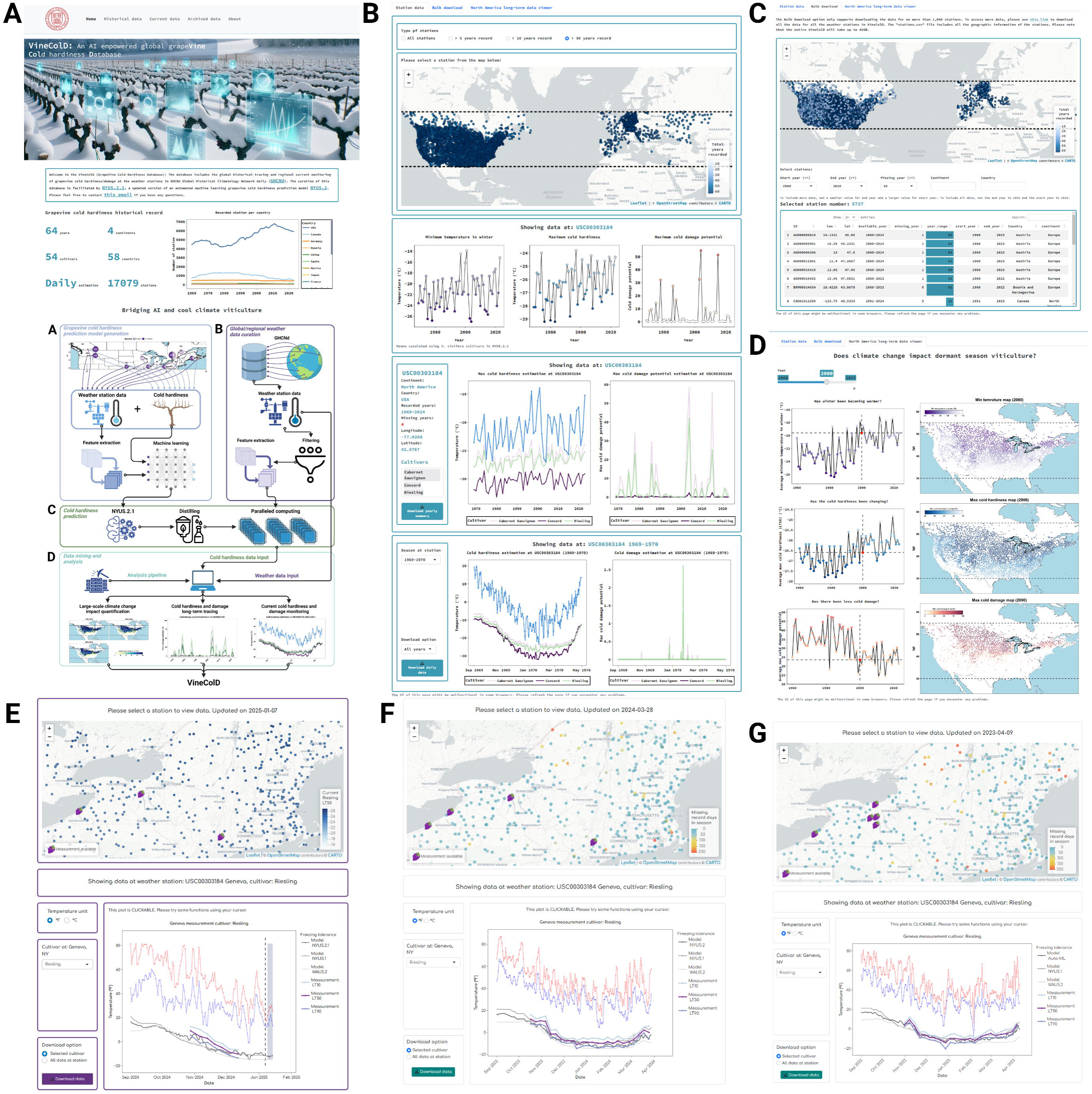
User interface of the R shiny-based application containing VineColD. A) Home page of the application; B) Composition of ‘Station data’ under ‘Historical data’ tab; C) Composition of ‘Bulk download’ under ‘Historical data’ tab; D) Composition of ‘North America long-term data viewer’ under ‘Historical data’ tab; E) Current user interface of the regional real-time grapevine cold hardiness monitoring application; F) Archived regional real-time grapevine cold hardiness monitoring application in the 2023-2024 dormant season; G) Archived regional real-time grapevine cold hardiness monitoring application in the 2022-2023 dormant season.

### Case study: using VineColD data to quantify the impact of climate change on grapevine cold damage in North America (1960 to 2024)

In our first attempt to utilize VineColD data, we conducted a case study analyzing grapevine cold hardiness and damage across North America using data from curated weather stations between 1960 and 2024. The goal was to identify trends and to compare grapevine cold hardiness and damage between different time periods to quantify the impact of climate change on cold damage during the dormant season. To facilitate this comparison, the period of 1960 to 2024 was segmented into three different time windows: 1960 to 1979, 1980 to 1999 and 2000 to 2024. For each window and at each curated weather station, we calculated the average minimum temperature, average maximum cold hardiness and average maximum cold damage during the dormant season. These calculations were performed for all cultivars, as well as specifically for V*. vinifera* and *V. hybrid* cultivars included in NYUS.2.1, to approximate the low-temperature boundaries for different grapevine species under cold stress. To enhance the spatial resolution of the analysis, we implemented a k-nearest neighbors (KNN) model using cold damage data from each window, with only longitude and latitude as input features [41]. This approach allowed us to extrapolate site-specific observations to a gridded spatial scale, resulting in the high-resolution maps presented in Figure 5.

**Figure 5.**
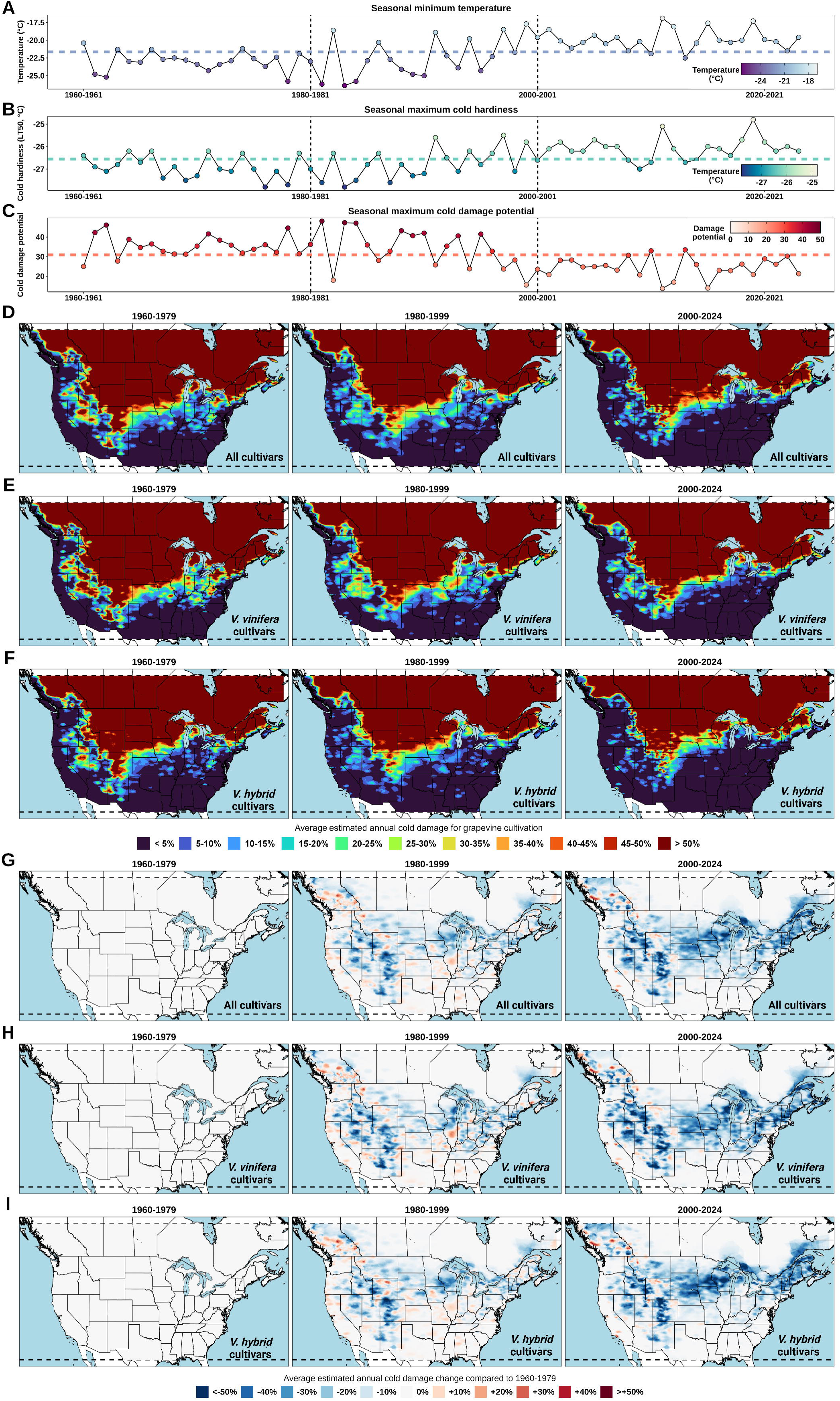
Quantification of the impact of climate change on grapevine cold damage in North America from 1960 to 2024. A) Average seasonally minimum temperature across all the curated weather stations. The line in the middle represents the average of the 64 dormant seasons; B) Average seasonally maximum estimated cold hardiness of all cultivars in NYUS.2.1 at all curated weather stations; C) Average seasonal maximum estimated cold damage potential of all cultivars in NYUS.2.1 at all curated weather stations; D) Gridded estimation of averaged seasonal cold damage of all cultivars included in NYUS.2.1 in North America, segmented into three time windows: 1960-1979, 1980-1999, and 2000-2024; E) Gridded estimation of averaged seasonal cold damage of *V. vinifera* cultivars included in NYUS.2.1 in North America between 1960 and 2024; F) Gridded estimation of averaged seasonal cold damage of *V. hybrid* cultivars included in NYUS.2.1 in North America between 1960 and 2024. G) Gridded estimation of averaged annual cold damage change of all cultivars compared to 1960-1979 H) Gridded estimation of averaged annual cold damage change of *V. vinifera* cultivars compared to 1960-1979; I) Gridded estimation of averaged annual cold damage change of *V. hybrid* compared to 1960-1979.

The regional analysis of estimated cold damage revealed significant shifts across North America from 1960 to 2024 (Figure 5). During the initial period of 1960-1979, average seasonal minimum temperatures at curated weather stations remained consistently low, ranging from -25.5 °C to - 20.4 °C (Figure 5A). Despite superior average seasonal maximum cold hardiness during this time, the average seasonal maximum cold damage potential was also high (Figures 5B and C). As a result, grapevine cultivation faced substantial cold damage risk, particularly in the upper Midwest, Great Lakes, and Northeast regions, with localized pockets of higher risk in the Rocky Mountain areas for *V. vinifera* (Figure 5E). For *V. hybrid* cultivars, the cold damage risk was more evenly distributed across the upper Midwest and Northeast, reflecting their broader cultivation potential and different cold hardiness characteristics (Figure 5F). In the 1980-1999 window, average seasonal minimum temperatures began to exhibit an “unstably increasing” trend, characterized by considerable fluctuations between years (Figure 5A). Similar fluctuating trends were observed in average seasonal maximum cold hardiness and cold damage potential (Figures 5B and C). Consequently, some small high-risk pockets emerged during this period in the regions such as the lower Midwest and South regions of the U.S. and British Columbia Province in Canada (Figure 5D to I). A regional decrease in cold damage risk was also observed in other regions such as the upper Midwest, Great Lakes, and Northeast regions of the U.S., as well as the lower regions of Ontario Province in Canada (Figures 5D to I). From 2000 to 2024, the trend of decreasing cold damage became more prevalent (Figures 5C to I). For *V. vinifera* cultivation, this decrease was evident in the Pacific Northwest, parts of the Midwest, and the Northeast, where cold damage risk fell below 20% in areas that previously faced higher risks (Figure 5E and H). Similarly, *V. hybrid* cultivars showed a notable decrease in cold damage risk in the Appalachian Highlands and coastal areas of the Pacific Northwest (Figure 5F and I). Notably, some coastal areas in the Maritime provinces of Canada and the New England states in the U.S. indicated potential for *V. hybrid* cultivation during this period (Figure 5F and I). Assuming regions with less than 20% annual cold damage are suitable for grapevine cultivation, our simulation predicts a 28.8% and 17.2% increase in the feasible land area for cultivating *V. vinifera* and *V. hybrid* cultivars, respectively, when comparing the 2000-2024 window to the 1960-1979 window.

## Conclusion

This study presents VineColD, a novel database that integrates historical and real-time data to evaluate grapevine cold hardiness, indicating a significant advancement in the modeling of grapevine physiology in dormant season. Utilizing the NYUS.2.1 model, VineColD enables precise, accurate and large-scale predictions of grapevine cold hardiness in the world, enhancing our understanding of grapevine physiology under diverse climatic conditions. To our knowledge, the current version of VineColD represents the first and the only large-scale grapevine cold hardiness database in the world. VineColD data estimates for the quantification of climate change’s impact on grapevine cold damage in North America indicate that there has been an increase in the land area feasible for grapevine cultivation if only considering cold damage as the limiting factor. Looking forward, the expansion VineColD coupled with the annual update of NYUS.2.1 to cover more cultivars and viticultural areas could provide a more comprehensive database for global grapevine cold hardiness/cold damage. VineColD sets a foundation for future research and development in viticulture, offering a robust platform for studying and mitigating the effects of climate change on grapevine cultivation. This innovative approach could lead to more resilient agricultural practices and contribute to the sustainability and growth of the viticulture industry worldwide.

## Data availability

VineColD is publicly available at https://cornell-tree-fruit-physiology.shinyapps.io/VineColD/. The current regional real-time grapevine cold hardiness monitoring system is publicly available at https://cornell-tree-fruit-physiology.shinyapps.io/North_America_Grape_Freezing_Tolerance/. The model training, model testing, feature importance analysis and model prediction are publicly available at https://github.com/imbaterry11/NYUS.2.

